# One-step multiple site-specific base editing by direct embryo injection for precision and pyramid pig breeding

**DOI:** 10.1101/2020.08.26.267948

**Authors:** Ruigao Song, Yu Wang, Qiantao Zheng, Jing Yao, Chunwei Cao, Yanfang Wang, Jianguo Zhao

## Abstract

Precise and simultaneous acquisition of multiple beneficial alleles in the genome to improve pig performance are pivotal for making elite breeders. Cytidine base editors (CBEs) have emerged as powerful tools for site-specific single nucleotide replacement. Here, we compare the editing efficiency of four CBEs in porcine embryonic cells and embryos to show that hA3A-BE3-Y130F and hA3A-eBE3-Y130F consistently results in higher base-editing efficiency and lower toxic effects to *in vitro* embryo development. We also show that zygote microinjection of hA3A-BE3-Y130F results in one-step generation of pigs (3BE pigs) harboring C-to-T point mutations, including a stop codon in *CD163* and in *MSTN* and induce beneficial allele in *IGF2*. The 3BE pigs showed improved growth performance, hip circumference, food conversion rate. Our results demonstrate that CBEs can mediate high throughput genome editing by direct embryo microinjection. Our approach allows immediate introduction of novel alleles for beneficial traits in transgene-free animals for pyramid breeding.

## Introduction

Innovative approaches to accelerate improvement of livestock is urgently needed to meet the increased demands for animal protein. Significant obstacles in livestock breeding are limited dispersion of beneficial traits between different species and time constrains required for crossbreeding and selection of livestock with improved performance. Genome editing techniques, such as the clustered regularly interspaced short palindromic repeat (CRISPR) system, have provided revolutionary progress for improvement of pig performance at reduced cost and shortened time (*Huang et al., 2014*; *Whitworth et al., 2014; Whitworth et al., 2016; Yang et al., 2011; Zhang et al., 2019*). CRISPR/Cas9 mediated knockout of the *MSTN* gene in pigs led to improved muscle development and decreased fat accumulation (*Wang et al., 2017*). Editing the regulatory element within intron 3 of the *IGF2* gene abolished repressor ZBED6 binding and resulted in improved meat production (*Xiang et al., 2018*). Most strikingly, studies of *CD163*-knockout pigs from the Prather group provided proof-of-concept that a single gene deletion establishes porcine reproductive and respiratory syndrome virus (PRRSV)-resistant pigs, which has been further confirmed by several researchers (*Burkard et al., 2018; Tanihara et al., 2019; Whitworth et al., 2016*). Most recently, base editors (BEs) that fuse a cytidine or adenosine deaminase with a catalytically impaired CRISPR–Cas9 mutant have been shown could directly and efficiently generate precise point mutations in genomic DNA, without generating double-strand breaks (DSBs) or requiring a donor template (*Gehrke et al., 2018*). These novel tools have been used to induce single nucleotide modifications in a variety of animals (*Kim et al., 2017; Liu et al., 2018; Xie et al., 2019*). Specifically, CBEs have been used to induce C-to-T conversions of multiple genes in cell clones and then used as donors in SCNT to make immune-deficient pigs (*Xie et al., 2019*). Thus, BEs provide a strategy for precise and efficient editing of single nucleotide polymorphisms (SNPs) to improve livestock traits in pigs.

As we know, SNPs are the richest and most abundant form of genomic polymorphisms that constitute the genetic architecture of economic traits (*Contreras-Soto et al., 2017; Mora et al., 2016; Yue et al., 2019*). Almost 66% of the total phenotypic variation of pigs can be explained by SNPs (*Lee et al., 2012*). Functional SNPs related to various economic traits such as meat quality and growth (*Lee et al., 2012*), fecundity (*Ma et al., 2018*) and virus resistance (*Hess et al., 2016*) have been identified. However, generation of gene-edited pigs harboring beneficial SNPs is very inefficient, has a low throughput and heavily relies on SCNT, a very complicated technology. The technical challenges of genetic engineering in somatic cells and SCNT greatly hinders widespread application of genome editing in livestock for breeding.

In the present study, we screened CBE variants and found hA3A-BE3-Y130F and hA3A-eBE3-Y130F consistently result in higher base-editing efficiency in PEFs and embryos. Pigs harboring C-to-T point mutations that create a stop codon in *CD163* and in *MSTN* and introduce a beneficial allele in intron 3–3072 of the *IGF2* gene were generated in one step with direct zygote microinjection of hA3A-BE3-Y130F. The genetically engineered pigs showed disrupted gene expression of *CD163* and *MSTN* and enhanced expression of *IGF2*, which resulted in improved growth performance. We found that CBEs mediated precise and efficient genome editing at multiple sites with direct embryo injection, circumventing the need for SCNT and providing a potential process for prospective pyramid breeding in pigs.

## Results

### CBEs-mediated base editing at multiple loci in PEFs

In order to screen suitable tools for base editing, the editing efficiency of four CBEs – rA1-BE3, hA3A-BE3, hA3A-BE3-Y130F and hA3A-eBE-Y130F – were compared in PEFs. Considering the potential for pyramid breeding, target loci from *CD163, MSTN* and *IGF2* genes were selected. Premature stop codons (R4312STOP and Q218STOP) were generated by a single C-to-T conversion at the target sites in *CD163* and *MSTN*, which are expected to confer double-muscled and porcine reproductive and respiratory syndrome virus (PRRSV) resistance traits in pigs, respectively (Fig. 1A). For *IGF2*, a point conversion at the target sites of Bama pig would abrogate BED-type containing 6 (ZBED6) binding, resulting in enhanced *IGF2* expression in skeletal muscle and improved lean meat yield (*Van Laere et al., 2003*) (Fig. 1A).

**Figure 1.**
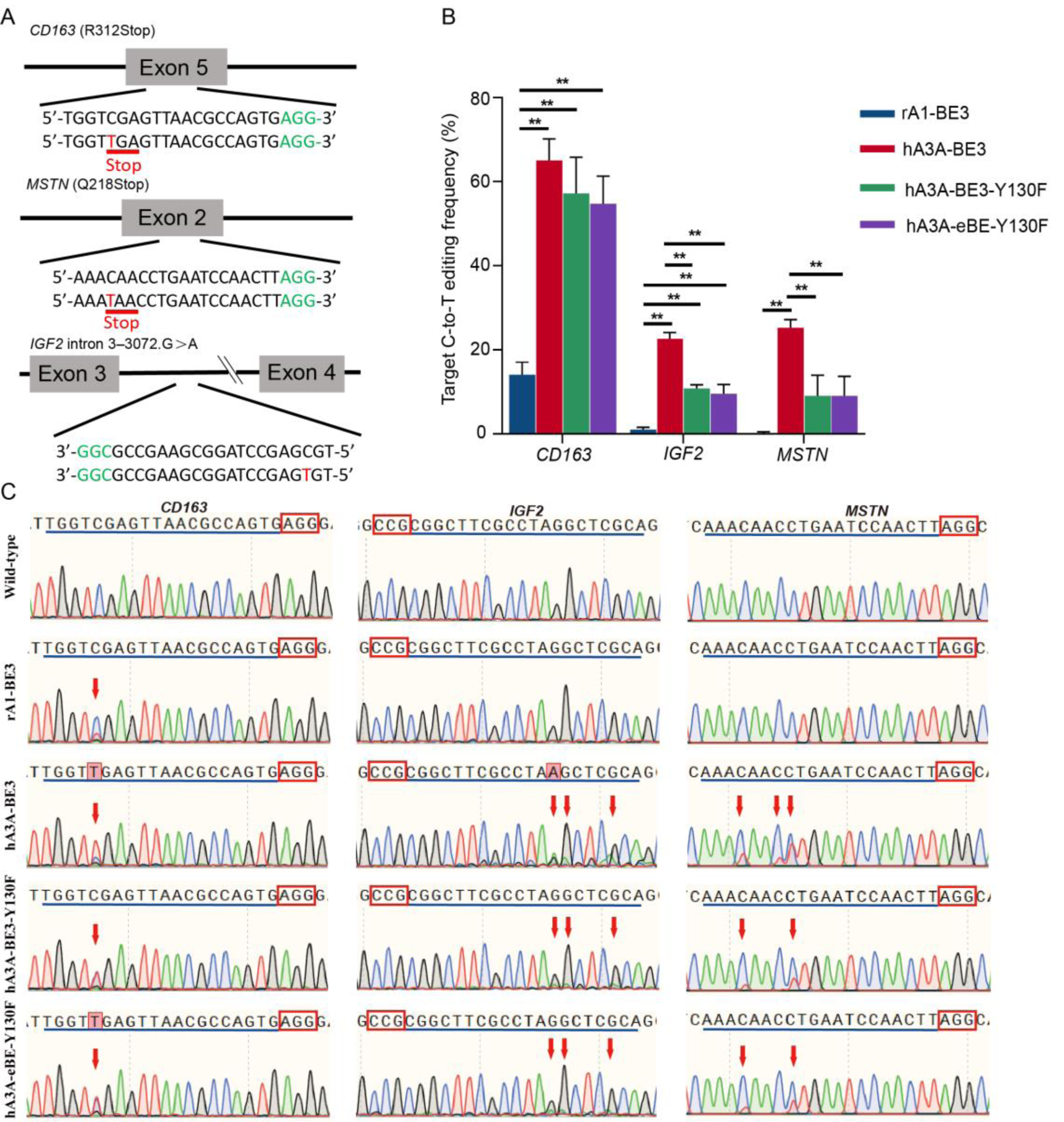
Base editing at multiple loci in PEF cells by different base editors. **A** Schematic of the *CD163, MSTN* and *IGF2* gene structure. One sgRNA targeting each gene was designed. The codon to be modified is underlined. The targeting site is in red and the PAM region is in green. **B** The C-to-T editing frequency of different base editors in multiple target sites were detected by sequencing. **C** Representative sequence chromatogram of different base editors in the target site of *CD163, MSTN* and *IGF2*. Red box shows the C-to-T substitutions at target sites.

Three sgRNA candidates targeting exons of porcine *CD163, MSTN* and *IGF2* genes were synthesized and transfected into PEFs with vectors expressing rA1-BE3, hA3A-BE3, hA3A-BE3-Y130F and hA3A-eBE-Y130F. After 48 h of culturing, cells were collected and target sequences were PCR amplified for sequencing. The editing efficiencies of hA3A-BE3, hA3A-BE3-Y130F and hA3A-eBE-Y130F were significantly higher than that of rA1-BE3 (Fig. 1B, C). We also observed the editing efficiencies varied among the three genes, with the *IGF2* target site showing lowest editing efficiency.

We next explored whether hA3A-BE3 could generate simultaneous C-to-T conversions in multiple genes in PEFs. The mixed sgRNAs were co-transfected with hA3A-BE3-expression vectors into PEFs, which were then cultured for 48 h before single-cell sorting by FACS. Next, thirty-nine single-cell colonies were selected and genotyped by Sanger sequencing after 8-14 days of cell culture. Among the 39 colonies, 28.21% (11/39) colonies showed C-to-T substitution among all three genes (Fig. 2A). In addition, 20 colonies (two for *CD163* and *IGF2*, sixteen for *CD163* and *MSTN*, and two for *IGF2* and *MSTN*) showed double-gene base editing, and 4 colonies (three for *CD163*, one for *MSTN*) showed single-gene base editing (Fig. 2A). Overall, 20.51% (8/39) colonies contained the targeted C-to-T homozygous mutation at all three gene sites (Fig. 2B). For each single gene, 76.92% (30/39), 28.21% (11/39) and 64.10% (25/39) of the colonies were homozygous mutants in *CD163, IGF2* and *MSTN*, respectively. Indel incorporation was also found in *CD163* (10.25%), *IGF2* (51.28%) and *MSTN* (15.38%) (Fig. 2C). In addition, an unwanted C-to-A substitution at the targeted position in *CD163* was detected in one colony (Fig. 2C). Thus, results show that hA3A-BE3 can efficiently induce biallelic C-to-T substitutions in three genes during one transfection of PEFs. Sanger sequence results further confirmed the high frequency of targeted C-to-T conversions (Fig. 2D).

**Figure 2.**
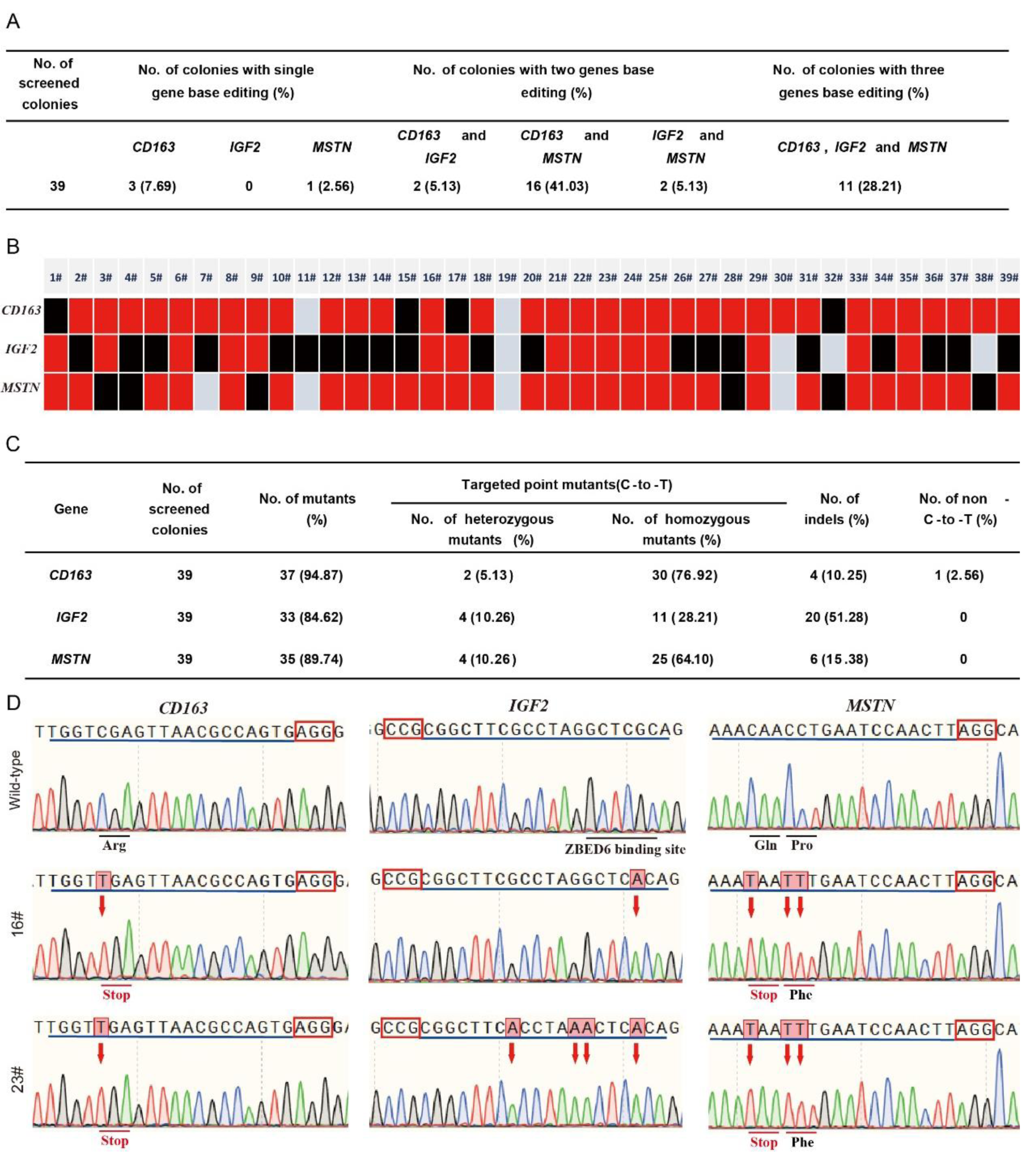
hA3A-BE3-mediated base editing for multiple genes in PEF cells. **A** Summary of base editing in PEF cells. **B** The base editing results of the *CD163, MSTN* and *IGF2* genes in the colonies. The red represents the base conversion of C-to-T, and the black represents an indels. **C** Summary of product types of base editing by hA3A-BE3 in porcine somatic cells. **D** Sanger sequencing chromatograms of the selected single-cell colonies. The red arrow indicate expected substitutions at target sites.

### Efficient one-step base editing for multiple genes in porcine embryos via embryo injection

We next detected the editing efficiency of these four CBEs in porcine embryos. In vitro transcribed sgRNAs (*CD163*-sgRNA and *MSTN*-sgRNA) and BE3, hA3A-BE3, hA3A-BE3-Y130F and hA3A-eBE-Y130F mRNAs were co-injected into porcine parthenogenetically activated (PA) oocytes. The injected PA embryos were cultured for 7 days to determine developmental competency. The blastocyst rate of embryos injected with *CD163*-sgRNA and mRNA of rA1-BE3, hA3A-BE3, hA3A-BE3-Y130F, and hA3A-eBE-Y130F were 15.8% (27/171), 0.96% (4/415), 15.63% (25/160) and 17.27% (24/139) (Fig. 3A), respectively. The blastocyst rate of embryos injected with *MSTN*-sgRNA and mRNA of rA1-BE3, hA3A-BE3, hA3A-BE3-Y130F, and hA3A-eBE-Y130F were 7.36% (12/163), 1.54% (9/585), 8.63% (22/255) and 8.52% (15/176) (Fig. 3B), respectively. The rates of embryos injected with CBEs are significantly lower than the rates of the control embryos injected with water, indicating that CBEs might affect early embryonic development competency. Notably, the hA3A-BE3 injection might lead to detrimental effects in embryo development, which may result from the high off-target rate of hA3A-BE3 in mRNA (*Zhou et al., 2019*).

**Figure 3.**
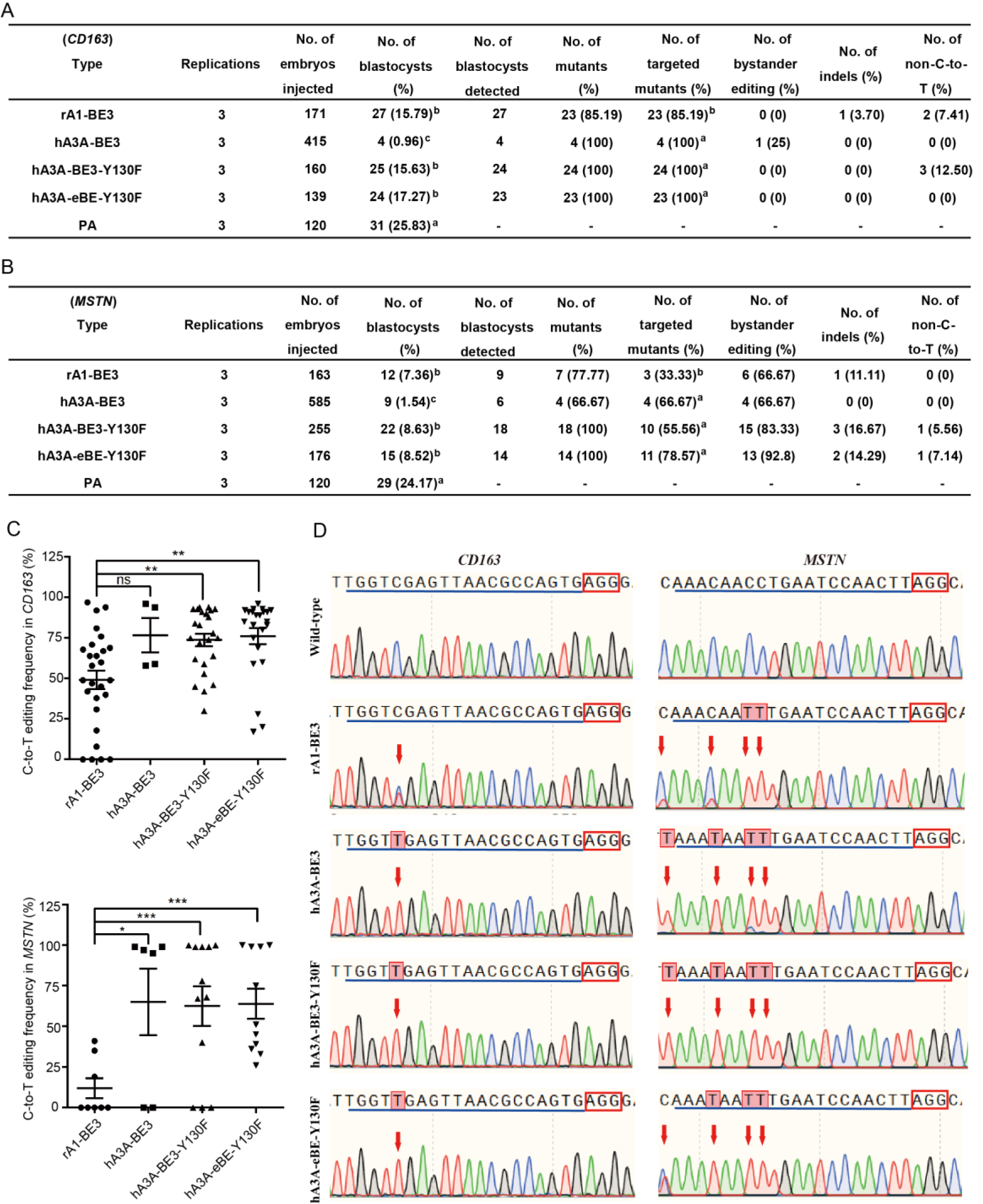
Efficient base editing for multiple sites in porcine embryos. Summary of base editing in *CD163* (**A**) and *MSTN* (**B**) by base editors in porcine embryos. **C** Summary of C-to-T editing frequency of base editors in *CD163*-C5 site and *MSTN*-C4 site in porcine embryos. **D** Sanger sequencing chromatograms of the different base editors in the *CD163* and *MSNT* genes in porcine embryos. The red arrow indicate substitutions at target sites. Superscript alphabets (a,b and c) represents values are significantly different (*P*<0.05).

Single blastocysts were lysed individually for genotyping by Sanger sequencing. Of the screened blastocysts, 85.2% (23/27) for rA1-BE3, 100% for hA3A-BE3 (4/4), 100% (24/24) for hA3A-BE3-Y130F and 100% (23/23) for hA3A-eBE-Y130F were identified to have targeted mutations in *CD163* (Fig. 3A). In one hA3A-BE3 injected blastocyst, C-to-T conversions also occurred outside the editing window. In addition to the specific C-to-T mutation, a few C-to-A substitutions were also found in rA1-BE3 (7.4%) and hA3A-BE3-Y130F (12.5%) injected blastocyst, as well as an indel in a rA1-BE3-injected blastocyst (Fig. 3A).

The targeted mutation rate in the *MSTN* injected group was 33.3% (3/9) for rA1-BE3, 66.7% (4/6) for hA3A-BE3, 55.6% (10/18) for hA3A-BE3-Y130F and 78.6% (11/14) for hA3A-eBE-Y130F (Fig. 3B). We also detected indels in 1 (11.1%, 1/9) blastocyst in the rA1-BE3 group, 3 (16.7%, 3/18) blastocysts in the hA3A-BE3-Y130F group and 2 (14.3%, 2/14) blastocysts in the hA3A-eBE-Y130F group (Fig. 3B). These data indicate rA1-BE3, hA3A-BE3-Y130F and hA3A-eBE-Y130F might induce DSB in the genome, resulting in indels (Fig. 3B). At the *MSTN* target site, we also have detected non-C-to-T conversion in 1 (5.6%, 1/18) blastocyst of the hA3A-BE3-Y130F group and 1 (8.3%, 1/12) blastocyst of the hA3A-eBE-Y130F group (Fig. 3B).

The targeted C-to-T mutation frequency in each blastocyst of different base editors is shown in Fig 3C and Fig 3D. At both the *CD163* and *MSTN* targeting sites, the editing efficiencies of hA3A-BE3-Y130F and hA3A-eBE-Y130F were significantly higher than that of rA1-BE3. In addition, the efficiency of hA3A-BE3 was significantly higher than that of rA1-BE3 at the *MSTN* gene site.

### Generation of *MSTN, CD163* and *IGF2* gene mutations in pigs via one-step zygote injection

Although hA3A-BE3 mediated the highest editing efficiency in both cell transfection and embryo injection, it also caused the greatest detrimental effect on embryonic development after injection. Hence, hA3A-BE3-Y130F was selected for pig zygote injection to base edit multiple genes in one step. The mixed sgRNAs and hA3A-BE3-Y130F mRNA was co-injected into the cytoplasm of one-cell stage zygotes obtained from nine Bama pigs. Thirty-eight injected zygotes were then transferred into three surrogate pigs (Fig. 4A). One surrogate pregnancy developed to term, resulting in a litter of five pigs with four healthy offspring (Fig. 4A, 4B). Genotyping showed that all five pigs harbored mutations in the three genes. At the target site of *CD163*, ∼100% of the site-specific C-to-T mutations was achieved at the *CD163* c.2142C site for all pigs, resulting in a stop codon with a 312 arginine conversion (Fig. 4C, D). The target deep sequence results showed that the intron 3-3072.G>A in the *IGF2* gene were detected in pigs with an efficiency of 46% (1920901), 98% (1920902) and 45% (1920905) (Fig. 4C, D). In addition, an unwanted mutation (intron 3-3067.G>A and intron 3-3068.G>A) were detected in pigs with an efficiency of 46% (1920901), 98% (1920902, 1920904) and 45% (1920905) (Fig. 4C, D). Moreover, three pigs (1920901, 1920904 and 1920905) harbored undesired indels at the *IGF2* target site (Fig. 4C, D). All five pigs harbored the target Q218Stop mutation in the *MSTN* gene, but two of the five pigs were heterozygous with a frequency of 50% and 41% (Fig. 4C, D). Unwanted L216L and P219F mutations were also detected in all pigs with high efficiency (Fig. 4C, D). Complete genotyping results are shown in Supplementary Fig. 1A and Fig. 1B. Notably, the 1920902 piglet harbored a biallelic C-to-T transition in three genes. Sequencing results indicate that all 3BE pigs were homozygous at *CD163* sites, one piglet (1920904) was homozygous at *MSTN* sites, and two pigs (1920902 and 1920903) were homozygous at *IGF2* sites (Fig 4D). In addition, we detected that 1920901, 1920902 and 1920905 were chimeric at *MSTN* sites. Sanger sequencing analyses of ten POTs showed that no off-target mutations were found in four of the base-edited pigs (Supplementary Fig. 2).

**Figure 4.**
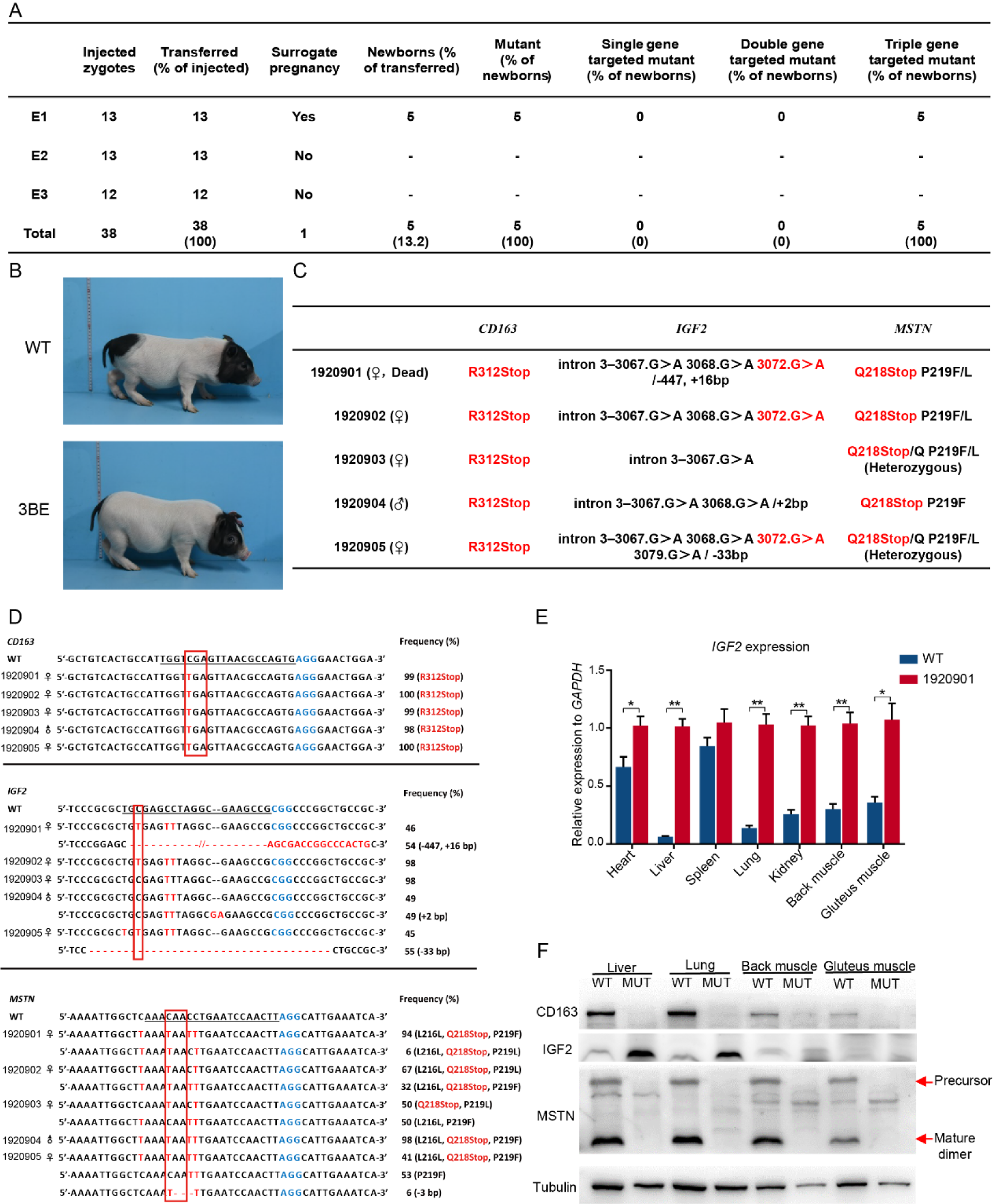
Generation *MSTN, CD163* and *IGF2* gene mutations in pigs via zygote injection. **A** Summary of generation of three genes mutant pigs by using direct zygote injection. **B** Representative photograph of newborn three genes mutant piglets. **C** The genotype of newborn piglets at *CD163, IGF2* and *MSNT*. **D** Summary of genotypes of newborn piglets from targeted deep sequencing. C-to-T substitutions and indels are shown in red. **E** Western blot was used to detect the expression of *CD163, IGF2* and *MSTN* protein in the liver, lung, back muscle and gluteus muscle of WT and 1920901 piglets. **F** The mRNA expression of *IGF2* in the heart, liver, spleen, lung, kidney, back muscle and gluteus muscle of WT and 1920901.

For mRNA expression analysis, q-PCR showed that the mRNA expression of *IGF2* in the heart, liver, lung, kidney, back muscle and gluteus muscle tissues of the piglet (1920901) was significantly higher than expression in wild-type (WT) pigs (Fig. 4E). The unregulated protein expression of IGF2 was also confirmed in the liver and lung of piglet (1920901) (Fig. 4F). Protein analysis demonstrated that CD163 and MSTN couldn’t be detected in the *CD163* R312Stop and *MSTN* Q218Stop mutant piglet (1920901) (Fig. 4F). However, the blood physiology and biochemistry reflected no significant difference between the 3BE and WT pigs (Table S2, S3).

### Improved growth performance in 3BE founder pigs

The body weight, body length, body height, hip circumference and chest circumference were measured in 3BE female pigs (n=3) and age matched WT female pigs (n=3). The birth weight of 3BE pigs was significantly higher than that of WT pigs (0.62±0.019 vs. 0.52±0.043, P=0.036) (Supplementary Fig.3 A). Results of growth performance showed that the 3BE pigs (0 – 6 months of age) grew faster than age-matched WT pigs (Fig.5 A). The hip circumference of 3BE pigs was also higher than that of WT pigs at every measuring time (Fig.5 B). The chest circumference, body length and body height of 3BE pigs were similar with WT pigs (Fig.5 C; Supplementary Fig.3 B, C). Results of average daily feed intake (ADF) (Fig.5 D), average daily gain (ADG) (Fig.5 E) and feed conversion ratio (FCR) showed that 3BE pigs (2 – 5 months of age) had lower FCRs than age-matched WT pigs (Fig.5 F). The pictures of 3BE pigs are shown in the Figure 5 G which indicate a higher hip circumference of 3BE pigs. Histological analysis of tissue slices of the gluteus muscle showed increased muscle fiber size in 3BE pigs compared with WT pigs (Fig. 5H, I).

**Figure 5.**
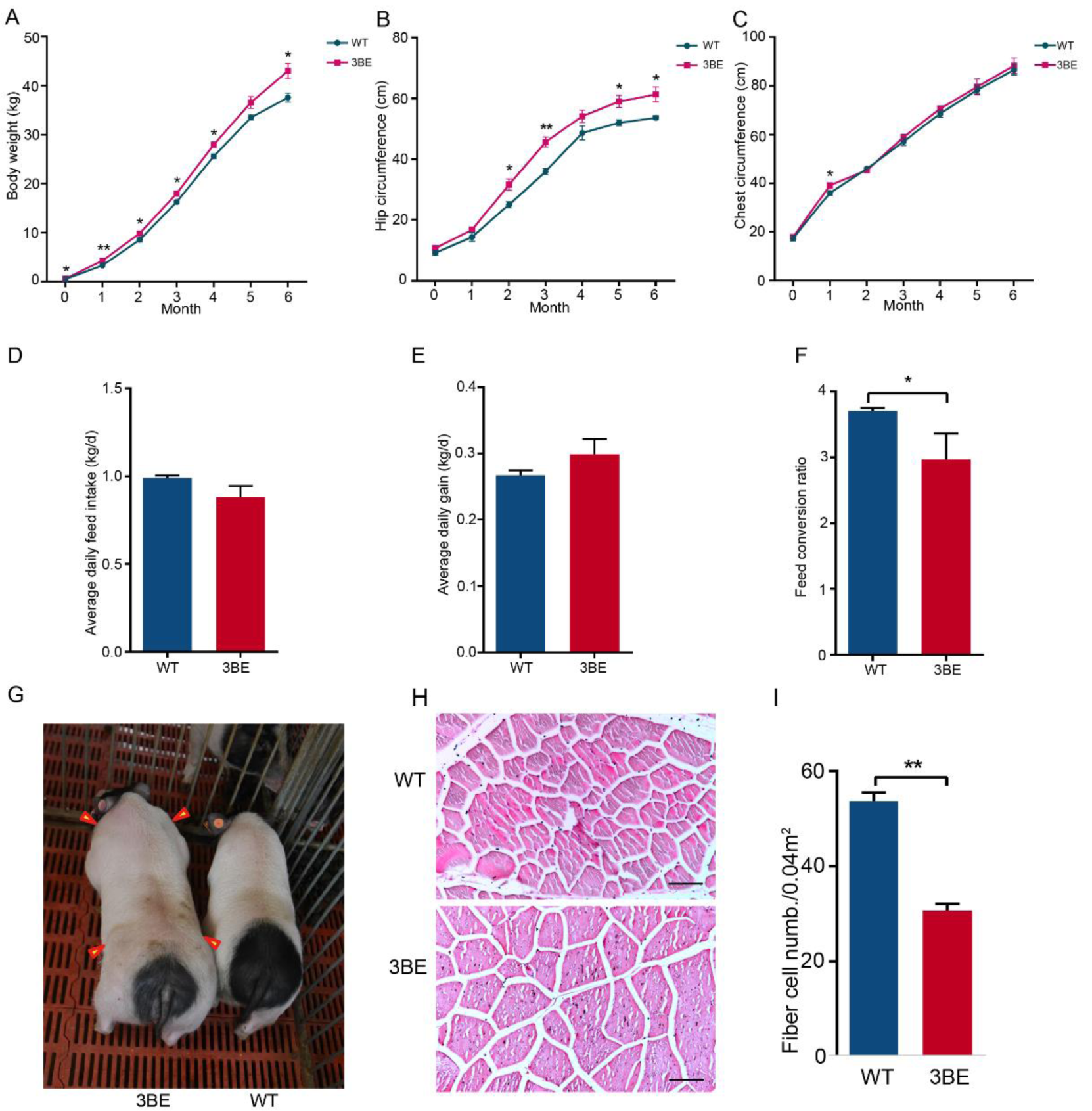
Growth performance of three genes mutant pigs. The body weight (**A**), saddle width (**B**), chest circumference (**C**), average daily feed intake (**D**), average daily gain (E) and feed conversion ratio (**F**) of 3BE pigs and age-matched WT (female and male) piglets were measured. **G** Images of 3 month-old female and male of 3BE and WT pig. **H, I** HE staining of gluteus muscle in WT and 3BE pigs and the changes of myofiber density in 3BE pigs.

## Discussion

Developing pig breeds with desirable characteristics such as high growth performance, meat quality, reproductive success, and disease resistance using traditional breeding methods is time consuming and cost prohibitive. Developing pigs using gene editing provides a reliable and rapid solution to improving livestock traits. In this study, a selective and efficient base editor was successfully used to engineer several beneficial traits by simultaneously editing multiply gene sites using one-step zygote injection in pigs.

With the continuous innovation and optimization of gene editing, many evolved versions of CBE tools have been developed to improve editing efficiency and accuracy (*Cheng et al., 2019; Koblan et al., 2018; Zong et al., 2018*). In this study, we first compared the editing efficiency of different CBEs at target sites in PEFs. All base editors were able to generate targeted site mutations, but the editing efficiency of CBEs varied at different target positions. The CBEs hA3A-BE3 and rA1-BE3 consistently showed the highest and lowest editing efficiency, respectively. In addition to the specific conversion of C-to-T mutation, undesired indels were also found at expected positions in these three genes. However, the efficiency of indel incorporation at the *IGF2* position was much higher than that of other genes. These results suggest that hA3A-BE3 may significantly improve the efficiency of base excision repair during C-to-T conversions in the high GC content regions (*Van Laere et al., 2003*) and subsequently induce indels in this region. These undesired results were in accordance with previous studies in human cells, which indicated the products may be dependent on uracil N-glycosylase (UNG) (*Komor et al., 2017*).

Even after the birth of Dolly more than two decades ago, SCNT is still quite inefficient, making the generation of genetically modified pigs through SCNT a continuous challenge. Zygote injection of Cas9 or BEs mRNA and sgRNA complexes to target single point mutation have been reported (*Liang et al., 2017; Liu et al., 2018*), but our current work is the first to characterize one-step multiple site-specific base editing. In this work, we found that almost all the injected CBE mRNA complexes have a negative influence on embryo development competence. This is especially true for hA3A-BE3 injections which may correspond with the high off-target rate of hA3A-BE3 in mRNA (*Zhou et al., 2019*). The editing efficiency of these CBE members were also variable depending on the differing targeted genes in embryos. Almost all embryos (85.2%∼100%) showed a single base mutation at the *CD163* target site, while 33.3%∼78.6% embryos showed a mutation at the *MSTN* target site. This variable editing efficiency may be associated with the different positions of the targeted site in genes. In addition, we demonstrated that the new CBEs (hA3A-BE3-Y130F and hA3A-eBE-Y130F) significantly increased targeted mutation efficiency in porcine blastocysts compared to their predecessor. However, at a cellular level, these new CBEs unexpectedly induced indels (0∼16.7%) and non-C-to-T conversions (0∼12.5%) in blastocysts at a higher frequency than that seen in a previous study (*Komor et al., 2016*). Other studies have also reported similar frequency of indels (6.7–35.6%) in *Psen1*-targeted mice (*Sasaguri et al., 2018*), and the different frequency of indels in genes were also found in the study of *Dmd*-targeted (11.1%) and *Tyr*-targeted (28.6%) mice (*Kim et al., 2017*).

Most economic traits in pigs are complex, quantitative, and usually controlled by multiple genes. Because of this complexity, trait improvement requires stacking of natural and induced mutations; therefore, precision and high-throughput single-base substitution will be vital for designed breeding. We share our proof-of-concept results that show CBE-mediated trait improvement in pigs is feasible. Using one-step zygote injection, we generated 3BE pigs that carried precise mutations in *CD163, IGF2* and *MSTN*. The hA3A-BE3-Y130F efficiently and precisely targeted C-to-T conversion with few proximal off-target mutations and negligible negative effects on embryo development, which is consistent with previous reports in mammalian cells (*Gehrke et al., 2018; St Martin et al., 2018*).

Our approach can help livestock breeders bypass time constraints with improving economic traits in pigs. Our approach can be used to directly create desirable mutations in a natural or genetically modified background using one-step zygote injection. To the best of our knowledge, this is the first report highlighting the highly efficient base conversion of three economic trait-related genes in pigs using a one step process. These findings indicate an achievable and rapid genetic improvement for multiple economic traits that conventional breeding and selection strategy could not accomplish. Beyond enhancing beneficial trait variation in pigs, our approach opens a new chapter in gene pyramid breeding for livestock.

## Materials and Methods

### Animals

Bama miniature pigs were maintained at the Beijing Farm Animal Research Center, Institute of Zoology, Chinese Academy of Sciences. All pig studies were conducted according to experimental practices and standards approved by the Institutional Animal Care and Use Committee of the Institute of Zoology, Chinese Academy of Sciences.

### Vector construction

*MSTN*-sgRNA, *CD163*-sgRNA and *IGF2*-sgRNA were designed following the NGG rule. Two complementary oligonucleotides of sgRNAs were synthesized and then annealed to double-stranded DNAs. The annealed products were then cloned into the BsaI-digested pGL3-U6-sgRNA expression vectors. sgRNA-oligo sequences used above are listed in Supplementary Table 1.

### mRNA and sgRNA preparation

rA1-BE3, hA3A-BE3, hA3A-BE3-Y130F and hA3A-eBE3-Y130F plasmids were obtained from Addgene. The plasmid was linearized with Not I, and mRNA was synthesized using an in vitro RNA transcription kit (mMESSAGE mMACHINE™ T7 ULTRA Transcription Kit, AM1345, Invitrogen). sgRNAs were amplified and transcribed in vitro using the MEGAshortscript™ T7 Transcription Kit (AM1354, Invitrogen) according to manufacturer’s instructions. sgRNA primer sequences are listed in Supplementary Table 1.

### PEFs culture and transfection

Pig fetal fibroblasts (PFFs) were isolated from 35-day-old fetuses of Bama pigs. A day before transfection, PFFs were thawed and cultured in PFFs culture medium (Dulbecco’s modified Eagle’s medium [DMEM, HyClone]) and supplemented with 15% fetal bovine serum (FBS, HyClone), 1% nonessential amino acids (NEAA, Gibco), and 2 mM GlutaMAX (Gibco). Next, ∼1 × 10^6^ PFFs were electroporated with rA1-BE3, hA3A-BE3, hA3A-BE3-Y130F or hA3A-eBE-Y130F (4 μg), and sgRNA-expressing vectors (2 μg of each sgRNA). The electroporated cells were recovered for 48 h, split into single cells and cultured in 96-well plates for 8 days to form colonies by flow cytometry. We also collected 10,000 single GFP positive cells to detect the efficiency of different base editor systems. Cell lysates were then used as templates for PCR. Next, PCR products were used for sub-cloning into the pMD18-T vector (Takara) and sequenced to determine mutation efficiency. Cell colonies with target mutations were identified using PCR and then sequenced. Primer sequences are listed in Supplementary table 1.

### Microinjection of pig embryo and zygotes

Porcine ovaries were obtained from a local slaughterhouse, and porcine oocyte collection, in vitro maturation, parthenogenetic activation (PA) and the reagent formulation were conducted as described in our previous studies (*Wang et al., 2015*). Zygotes from Bama miniature pigs were collected within 24 h after insemination and transferred into manipulation medium (0.75 g HEPES, 9.5 g TCM-199 powder, 0.05 g NaHCO_3_, 1.755 g NaCl, 0.05 g penicillin, 0.06 g streptomycin, and 3.0 g BSA, in a final volume of 1L in Milli-Q water, pH 7.2–7.4). The protocol for microinjection of oocytes and pronuclear stage embryos has been described in detail in our published protocols (*Wang et al., 2015*). Briefly, a mixture of rA1-BE3, hA3A-BE3, hA3A-BE3-Y130F and hA3A-eBE3-Y130F mRNA (200 ng/ul), and sgRNA (50 ng/ul) was co-injected into the cytoplasm of porcine PA embryos and pronuclear-stage zygotes. The injected embryos or zygotes were then cultured in PZM3medium, pH 7.4, supplemented with 3 mg/ml BSA, for 7 days or transferred to surrogate pigs.

### Single embryo PCR amplification and pig genotyping

Injected embryos were collected at the blastocyst stage. Genomic DNA was extracted with lysis buffer (1% NP40) at 55 °C for 40 min and 95 °C for 15 min, and then were used as a template for PCR, and subjected to Sanger sequencing. Genomic DNA of newborn pigs was extracted from ear clips for PCR and subjected to Sanger sequencing and targeted deep sequencing. All primers for detection are listed in Supplementary Table 1.

### Targeted deep sequencing

Target sites were amplified from genomic DNA using i5 Phusion polymerase. The paired-end sequencing of PCR amplicons was performed by Sangon Biotech (Shanghai), using an Illumina MiSeq. The primers are listed in supplementary Table 1.

### Quantitative real-time PCR

Total RNA was extracted from porcine heart, liver, spleen, lung, kidney, back muscle and gluteus muscle tissues with TRIzol (Invitrogen). Total RNAs were used for reverse transcription using a FastQuant RT Kit (Tiangen Biotech). qPCR reactions were performed using TaKaRa SYBR Premix Ex Taq (TaKaRa, Tokyo, Japan) with an Agilent Mx3005p (Agilent Technologies) quantitative PCR instrument. The housekeeping gene, *GAPDH*, was used as an internal control. Relative expression of *IGF2* was calculated using the comparative cycle threshold (2^-ΔΔCt^) method. All primer sequences are shown in Supplementary Table 1.

### Western blot analysis

Liver, lung, back muscle and gluteus muscle tissues were dissected and frozen immediately in liquid nitrogen and stored at -80°C until use. Total protein was extracted using the Minute Total Protein Extraction Kit (Invent Biotechnologies, Inc.). Proteins were loaded to SDS-PAGE and transferred onto PVDF membranes. Membranes were blocked with 5% fat-free milk for 1 h at room temperature. Primary antibodies, anti-CD163 (ab87099, 1:500; Abcam), anti-IGF2 (ab170304, 1:100; Abcam) anti-MSTN (ab98337, 1:100; Abcam), and anti-Tubulin (SC9104, 1:3,000; Santa Cruz) were incubated with the membranes overnight at 4°C. Next, membranes were incubated with HRP-conjugated secondary antibodies for 1h at room temperature. All signals were detected using ECL Prime chemiluminescence (GE Healthcare) according to the manufacturer’s protocols.

### Off-target assay

Ten potential off-target sites (POTs) for each sgRNA were predicted for site-specific edits by a base-editor system, according to an online design tool CAS-OFFinder (http://www.rgenome.net/cas-offinder/)(*Bae et al., 2014*). All POTs were amplified by PCR and then subjected to sequencing. All primers for amplifying the off-target sites are listed in Supplementary table S1.

### Histochemistry

Three tissue samples (0.5 cm × 0.5 cm × 0.5 cm) were randomly chosen from the gluteus muscles of 3BE pigs at 6 months of age. The tissues were fixed in 4% paraformaldehyde for 24 h and then moved into 70% alcohol. Samples were embedded in wax blocks, sectioned at 5μm, and then stained with hematoxylin and eosin (H&E). Five different microscopic fields were randomly chosen, and muscle fiber density was defined as the number of myofibers per 0.04 mm^2^ of the muscle cross-sectional area.

### Blood analysis

Venous blood samples were collected to determine blood physiology and biochemistry. Samples were tested in the Beijing Lawke Health Laboratory.

## Statistical analysis

Statistical data are expressed as mean ± SEM and at least three individual representations were included in all experiments. Statistical significance was analyzed with a Student’s t test (unpaired) or two-way ANOVA using GraphPad prism software 6.0. A p-value < 0.05 was considered statistically significant. **, *P* < 0.01; *, *P* < 0.05.

## Acknowledgement

This work was supported by the National Natural Science Foundation for distinguished Young Scholars (31672387), National Key Research and Development Program of China (2020YFC1316602); National Natural Science Foundation of China (81671274, 31925036, 31272440, and 31801031), the National Transgenic Project of China (2016ZX08009003-006-007), and the Elite Youth Program of the Chinese Academy of Agricultural Sciences (ASTIP-IAS05).

## Author contributions

Jianguo Zhao and Yanfang Wang conceived the study. Ruigao Song, Yu Wang and Qiantao Zheng designed and performed the experiments and analyzed the data. Ruigao Song, Yu Wang performed molecular experiments. Ruigao Song performed embryo microinjection and embryos transfer experiments. Cunwei Cao analyzed the off-target date. Ruigao Song wrote the manuscript. Yanfang Wang and Jianguo Zhao supervised the project and revised the paper. All authors read and approved the final manuscript.

Conflict of interest

The authors declare that they have no conflict of interest.

## Notes

### Competing Interest Statement

The authors have declared no competing interest.

