## Supplemental figure1-3 and supplemental Table1-3 for "One-step multiple site-specific base editing by direct embryo injection for precision and pyramid pig breeding"

### Supplemental files

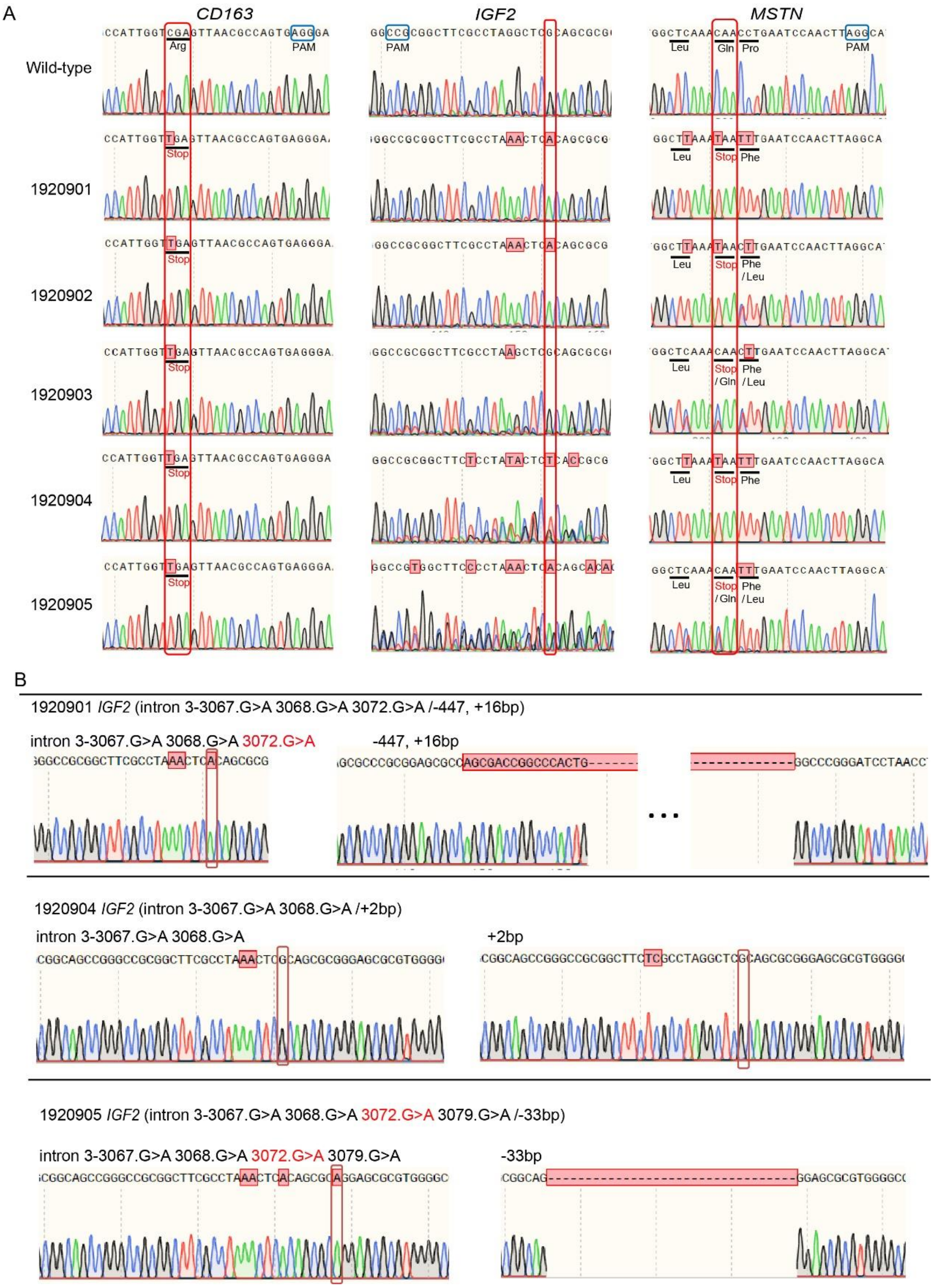


**Figure S1 Genotyping of 3BE pigs via Sanger sequencing. A** Sanger sequencing chromatograms of DNA from five cloned piglets and WT. The red arrow indicates the substituted nucleotides. **B** Sanger sequencing of 1920901, 1920904 and 1920905 piglets in the *IGF2* gene.

A

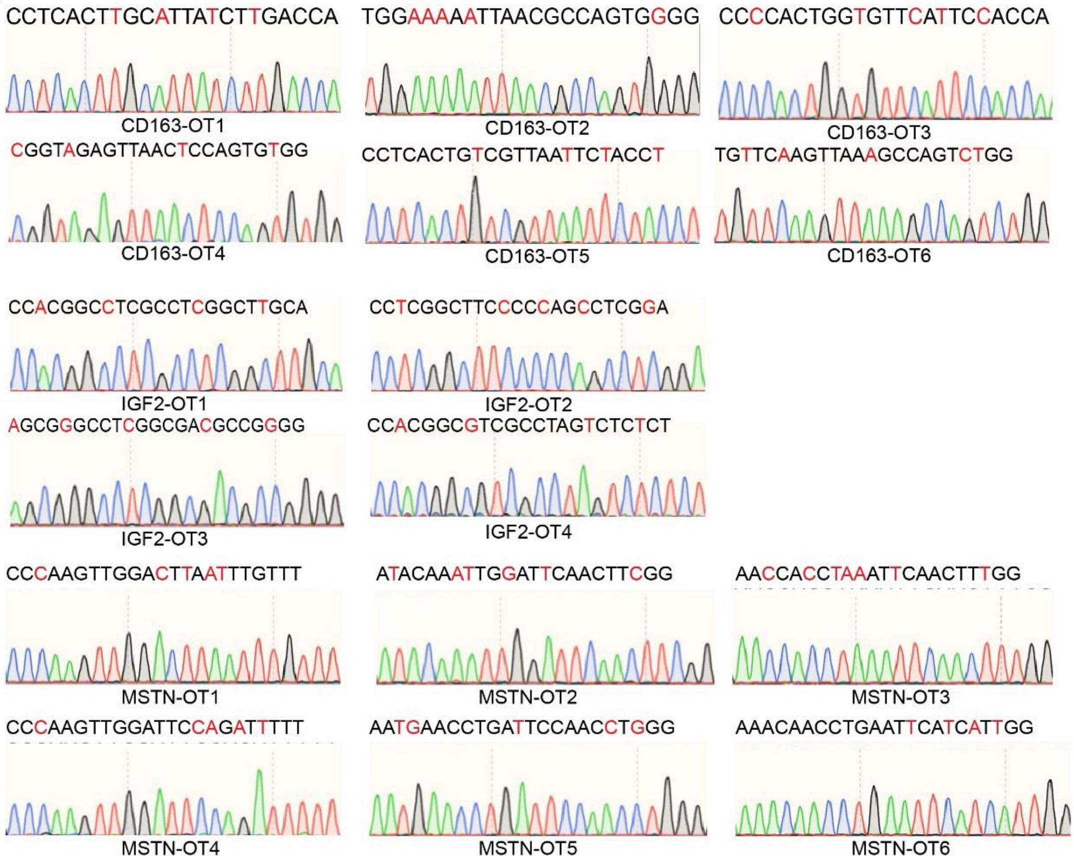


B

| | Sequences | Position in genome | Results |
|---|---|---|---|
| CD163 Target sites | TGGTCGAGTTAACGCCAGTGAGG | | |
| CD163 OT-1 | CCTCACTTGCATTATCTTGACCA | NC_010448.4_67498434 | No-off-target occurred |
| CD163 OT-2 | TGGAAAAATTAACGCCAGTGGGG | NC_010458.4_72856132 | No-off-target occurred |
| CD163 OT-3 | CCCCACTGGTGTTCATTCCACCA | NC_010455.5_142058382 | No-off-target occurred |
| CD163 OT-4 | CGGTAGAGTTAACTCCAGTGTGG | NC_010456.5_58117197 | No-off-target occurred |
| CD163 OT-5 | CCTCACTGTCGTTAATTCTACCT | NC_010452.4_56447489 | No-off-target occurred |
| CD163 OT-6 | TGTTCAAGTTAAAGCCAGTCTGG | NC_010453.5_10546882 | No-off-target occurred |
| IGF2 Target sites | TGCGAGCCTAGGCGAAGCCGCGG | | |
| IGF2 OT-1 | CCACGGCCTCGCCTCGGCTTGCA | NC_010448.4_64326215 | No-off-target occurred |
| IGF2 OT-2 | CCTCGGCTTCCCCCAGCCTCGGA | NC_010444.4_37255067 | No-off-target occurred |
| IGF2 OT-3 | AGCGGGCCTCGGCGACGCCGGG[illegible] | NC_010446.5_84204845 | No-off-target occurred |
| IGF2 OT-4 | CCACGGCGTCGCCTAGTCTCTCT | NC_010447.5_61425713 | No-off-target occurred |
| MSTN Target sites | AAACAACCTGAATCCAACTTAGG | | |
| MSTN OT-1 | CCCAAGTTGGACTTAATTTGTTT | NC_010456.5_66056237 | No-off-target occurred |
| MSTN OT-2 | ATACAAATTGGATTCAACTTCGG | NC_010447.5_91280658 | No-off-target occurred |
| MSTN OT-3 | AACCACCTAAATTCAACTTTGG | NC_010459.5_42354543 | No-off-target occurred |
| MSTN OT-4 | CCCAAGTTGGATTCCAGATTTTT | NC_010454.4_27795172 | No-off-target occurred |
| MSTN OT-5 | AATGAACCTGATTCCAACCTGGG | NC_010461.5_91265948 | No-off-target occurred |
| MSTN OT-6 | AAACAACCTGAATTCATCATTGG | NC_010455.5_77147926 | No-off-target occurred |

**Figure S2. No off-target site was detected in the piglets. A** Sanger sequencing of the potential targeting sites. **B** The target sequences and their genome positions.

A

| groups | Birth weight (kg) |
|---|---|
| WT | 0.52 ± 0.056 [b] |
| 3BE | 0.62 ± 0.020 [a] |

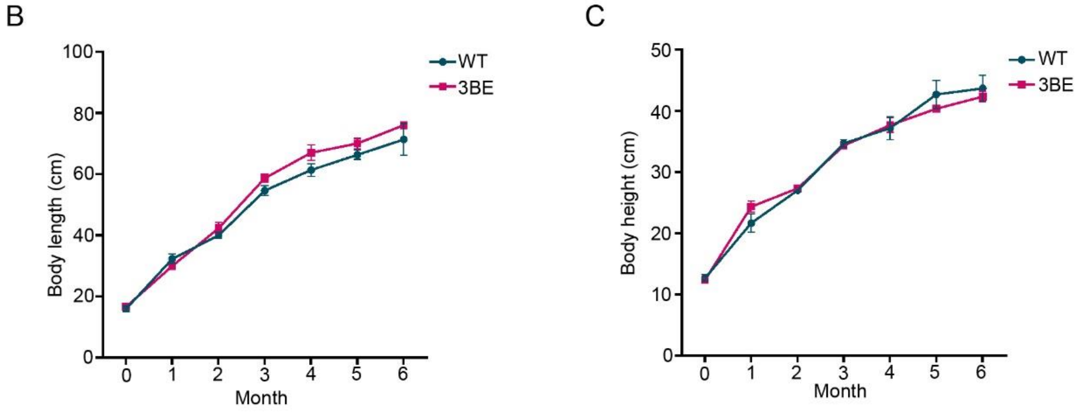


**Figure S3** The body length and height of 3BE pigs were similar with WT pigs. **A** The birth weight of 3BE pigs. The body length (**B**) and body height (**C**) of 3BE and age-match WT pigs.

**S1 Table 1. Oligonucleotides used in this study.**

Oligonucleotides for gRNA

| Primer name | Sequence |
|---|---|
| *CD163* gRNA 1F | CACCGTGGTCGAGTTAACGCCAGTG |
| *CD163* gRNA 1R | AAAC CACTGGCGTTAACTCGACCA |
| *IGF2* gRNA 2F | CACCGTGCGAGCCTAGGCGAAGCCG |
| *IGF2* gRNA 2R | AAACCGGCTTCGCCTAGGCTCGCA |
| *MSTN* gRNA 3F | CACCGAAACAACCTGAATCCAACTT |
| *MSTN* gRNA 3R | AAACAAGTTGGATTCAGGTTGTTT |

Oligonucleotides for genotyping

| Primer name | Sequence |
|---|---|
| *pCD163-GT-E5-F2* | TCCCCTTTTCACTCACTCCTC |
| *pCD163-GT-E5-R2* | GGTCCTTCCGGTGTTTTGTTTTC |
| *pIGF2-GT-F9* | CACTCTCTCAATTCCCCAAGCA |
| *pIGF2-GT-R9* | TAGTGCCCGAAAAACCCACTT |
| *pMSTN-GT-E2-F1* | TGGATGTTCCTCCACAGTGTC |
| *pMSTN-GT-E2-R1* | CAGGGCTACCATTGGGGTAA |

Oligonucleotides for real-time PCR

| Primer name | Sequence |
|---|---|

| | |
|---|---|
| *pIGF2-RT-F1* | GTGCTGCTATGCTGCTTACCG |
| *pIGF2-RT-R1* | CCGCAGACAAACTGGAGGG |
| *pGAPDH-RT-F1* | CCTTCATTGACCTCCACTACATGGT |
| *pGAPDH-RT-R1* | CCACAACATACGTAGCACCAGCATC |

#### Oligonucleotides for in vitro transcription

| Primer name | Sequence |
|---|---|
| pMSTN-NG-g3-IV-F | TAATACGACTCACTATAGGaaacaacctgaatccaacttGTTTTAGAGCTAGAAATAGC |
| pIGF2-I3-sgRNA-IV-F | TAATACGACTCACTATAGGtgCgagcctaggcgaagccgGTTTTAGAGCTAGAAATAGC |
| pCD163-E5-sgRNA-IV-F | TAATACGACTCACTATAGGtggtcgagttaacgccagtgGTTTTAGAGCTAGAAATAGC |
| In vitro transcript-Rev primer | TAATGCCAACTTTGTACAAGAAAG |

#### Primer used for targeted deep sequence

| Primer name | Sequence |
|---|---|
| CD163-P1-TS-F | ATCACGACAGCTCTTCAGTTTGCCCT |
| IGF2-P1-TS-F | ATCACGTCCTTTCCCAGTCCTTCCAC |
| MSTN-P1-TS-F | ATCACGGACCCGTCAAGACTCCTACA |
| CD163-P1-TS-R | CGATGTCCAGAACATGTCACACCAGC |
| IGF2-P1-TS-R | TTAGGCCGCTTCTCCTGCCACTGA |
| MSTN-P1-TS-R | TGACCAACTTTTATTGGGTACAGGGCTAC |
| CD163-P2-TS-F | ACAGTGACAGCTCTTCAGTTTGCCCT |
| IGF2-P2-TS-F | ACAGTGTCCTTTCCCAGTCCTTCCAC |
| MSTN-P2-TS-F | ACAGTGGACCCGTCAAGACTCCTACA |
| CD163-P2-TS-R | GCCAATCCAGAACATGTCACACCAGC |
| IGF2-P2-TS-R | CAGATCCGCTTCTCCTGCCACTGA |
| MSTN-P2-TS-R | ACTTGAACTTTTATTGGGTACAGGGCTAC |
| CD163-P3-TS-F | GATCAGACAGCTCTTCAGTTTGCCCT |
| IGF2-P3-TS-F | GATCAGTCCTTTCCCAGTCCTTCCAC |
| MSTN-P3-TS-F | GATCAGGACCCGTCAAGACTCCTACA |
| CD163-P3-TS-R | TAGCTTCCAGAACATGTCACACCAGC |
| IGF2-P3-TS-R | GGCTACCGCTTCTCCTGCCACTGA |
| MSTN-P3-TS-R | CTTGTAACTTTTATTGGGTACAGGGCTAC |

| | |
|---|---|
| CD163-P4-TS-F | AGTCAAACAGCTCTTCAGTTTGCCCT |
| IGF2-P4-TS-F | AGTCAATCCTTTCCCAGTCCTTCCAC |
| MSTN-P4-TS-F | AGTCAAGACCCGTCAAGACTCCTACA |
| CD163-P4-TS-R | AGTTCCCCAGAACATGTCACACCAGC |
| IGF2-P4-TS-R | ATGTCACGCTTCTCCTGCCACTGA |
| MSTN-P4-TS-R | CCGTCCACTTTTATTGGGTACAGGGCTAC |
| CD163-P5-TS-F | GTAGAGACAGCTCTTCAGTTTGCCCT |
| IGF2-P5-TS-F | GTAGAGTCCTTTCCCAGTCCTTCCAC |
| MSTN-P5-TS-F | GTAGAGGACCCGTCAAGACTCCTACA |
| CD163-P5-TS-R | GTCCGCCCAGAACATGTCACACCAGC |
| IGF2-P5-TS-R | GTGAAACGCTTCTCCTGCCACTGA |
| MSTN-P5-TS-R | GTGGCCACTTTTATTGGGTACAGGGCTAC |

#### Oligonucleotides for putative off-target sites

| Primer name | Sequence |
|---|---|
| pCD163-off-target-1-F1 | GCCTCCCTGTGTCAACATCA |
| pCD163-off-target-1-R1 | CAGTGTTTGTGCAGACGCTC |
| pCD163-off-target-2-F1 | TCCTGTGGTTTTGTCGTGCA |
| pCD163-off-target-2-R1 | AGCCAGGGAGGTGAGAGTAG |
| pCD163-off-target-3-F1 | GAGGGGTGGGGATTTGACTG |
| pCD163-off-target-3-R1 | TGGTCAGTGTGGCTGTTTGA |
| pCD163-off-target-4-F1 | CAAACCCGGGAGAACTCAGG |
| pCD163-off-target-4-R1 | ATCACCCAGGAGGAAGCTCT |
| pCD163-off-target-5-F1 | AGAGTTTGAGTCACTGGCGG |
| pCD163-off-target-5-R1 | ACTGCCACAGAGTTTGCAGT |
| pCD163-off-target-6-F1 | ACAGGTGTAGAGCAGGTAGACT |
| pCD163-off-target-6-R1 | TGAGCTGTCTACATAAAGAGGGA |
| pIGF2-off-target-1-F1 | GGAAGTGAGGCCTCTGTCAG |
| pIGF2-off-target-1-R1 | CTCTGGGATCAGCAGCAGAC |
| pIGF2-off-target-2-F1 | CGCTGGAGCTGAGGAAGAAA |
| pIGF2-off-target-2-R1 | GTTCCGGGAAAGGTACTGGG |
| pIGF2-off-target-3-F1 | CATCCTTTGCGTCTCTCCGG |
| pIGF2-off-target-3-R1 | GAAAGATCCCAGACCCGTCC |
| pIGF2-off-target-4-F1 | GATACCTCACAGGCCTGCTG |
| pIGF2-off-target-4-R1 | CGTGTCCAAACTTGCCCTTG |
| pMSTN-off-target-1-F1 | CCCTACTTCTGGGTTCACTGC |
| pMSTN-off-target-1-R1 | TTTTCACCAGTGGAGCACCA |
| pMSTN-off-target-2-F1 | GATGGCAGCAAACATCCACC |

| | |
|---|---|
| pMSTN-off-target-2-R1 | TTTCTGGTGGTGAAGGGTGG |
| pMSTN-off-target-3-F1 | TGAGGAACAAACCCCAGAGG |
| pMSTN-off-target-3-R1 | ACTCCTCGAGGGATCTTTCA |
| pMSTN-off-target-4-F1 | AGTTATGCACAGCACCCACA |
| pMSTN-off-target-4-R1 | CCTCCAGTGGGCATCTAAGC |
| pMSTN-off-target-5-F1 | CCTAGGGGCTCTGCTTTGTC |
| pMSTN-off-target-5-R1 | GGAGGAATCCCTGTGCCAAA |
| pMSTN-off-target-6-F1 | CACAAGGAAGCGGAGAGTGT |
| pMSTN-off-target-6-R1 | CTTCTGGGCTCAGGGAGTTG |

#### Table S2. Blood routine examination of 3BE and WT pigs.

| Items | WT | | 3BE | | P-value |
|---|---|---|---|---|---|
| | n | Mean ± S.E.M. | n | Mean ± S.E.M. | |
| WBC ($*10^9$/L) | 3 | 16.15 ± 2.53 | 3 | 14.44 ± 4.83 | 0.24 |
| RBC ($*10^{12}$/L) | 3 | 8.23 ± 0.29 | 3 | 9.44 ± 2.87 | 0.07 |
| HGB (g/L) | 3 | 160.67 ± 7.72 | 3 | 161.67 ± 51.23 | 0.73 |
| HCT (%) | 3 | 49.32 ± 2.39 | 3 | 48.99 ± 15.61 | 0.92 |
| MCV (fL) | 3 | 54.90 ± 7.02 | 3 | 51.97 ± 16.15 | 0.87 |
| MCH (pg) | 3 | 18.20 ± 1.91 | 3 | 17.14 ± 5.43 | 0.85 |
| MCHC (g/L) | 3 | 329.67 ± 3.77 | 3 | 329.67 ± 107.82 | 0.65 |
| RDW | 3 | 23.03 ± 4.36 | 3 | 28.37 ± 8.96 | 0.94 |
| PLT ($*10^9$/L) | 3 | 299.33 ± 42.30 | 3 | 266.67 ± 96.34 | 0.14 |
| MPV | 3 | 9.10 ± 0.93 | 3 | 9.70 ± 2.89 | 0.88 |
| PDW | 3 | 17.50 ± 0.22 | 3 | 18.03 ± 5.80 | 0.10 |
| NE% | 3 | 19.13 ± 6.77 | 3 | 22.00 ± 6.85 | 0.82 |
| LY% | 3 | 68.97 ± 26.24 | 3 | 71.40 ± 21.99 | 0.93 |
| MO% | 3 | 11.32 ± 5.28 | 3 | 10.29 ± 3.91 | 0.68 |
| EO% | 3 | 0.26 ± 0.03 | 3 | 0.19 ± 0.12 | 0.80 |
| BA% | 3 | 0.51 ± 0.44 | 3 | 0.61 ± 0.28 | 0.75 |

**WBC,** white blood cell count; **RBC,** red blood cell count; **HGB**, hemoglobin; **HCT**, hematocrit; **MCV**, mean corpuscular volume; **MCH**, mean corpuscular hemoglobin; **MCHC**, mean corpuscular hemoglobin concentration; **RDW,** red cell volume distribution width; **PLT**, platelet; **MPV**, mean platelet volume; **PDW**, platelet distributing width; **NE%**, neutrophilic granulocyte percentage; **LY%**, Lymphocyte percentage; **MO%**, monocytes percentage; **EO%**, eosinophilic cells percentage; **BA%**, basophils percentage. Quantitative data are presented as Mean ± S.E.M. Significance was established using Student's t-test. Differences were considered significant at *P<0.05.

Table S3. Blood biochemical examination of 3BE and WT pigs.

| Items | WT | | 3BE | | P-value |
|---|---|---|---|---|---|
| | n | Mean ± S.E.M. | n | Mean ± S.E.M. | |
| ALT | 3 | 33.60 ± 3.06 | 3 | 29.90 ± 9.94 | 0.19 |
| AST | 3 | 43.70 ± 13.70 | 3 | 70.63 ± 21.64 | 0.10 |
| GGT | 3 | 54.23 ± 1.73 | 3 | 56.93 ± 18.28 | 0.61 |
| ALP | 3 | 112.93 ± 53.09 | 3 | 87.17 ± 38.92 | 0.54 |
| AMY | 3 | 2770.67 ± | 3 | 2577.33 ± | 0.81 |
| TP | 3 | 75.33 ± 4.81 | 3 | 79.57 ± 24.41 | 0.45 |
| ALB | 3 | 40.37 ± 0.68 | 3 | 36.05 ± 12.87 | 0.17 |
| Mg | 3 | 0.78 ± 0.02 | 3 | 0.75 ± 0.25 | 0.63 |
| TBIL | 3 | 0.80 ± 0.09 | 3 | 0.95 ± 0.27 | 0.09 |
| TBA | 3 | 7.73 ± 3.74 | 3 | 9.73 ± 3.44 | 0.58 |
| $TCO_2$ | 3 | 20.57 ± 4.39 | 3 | 19.30 ± 5.89 | 0.71 |
| Urea | 3 | 2.02 ± 0.34 | 3 | 2.61 ± 0.77 | 0.20 |
| TC | 3 | 12.20 ± 7.31 | 3 | 1.90 ± 6.55 | 0.12 |
| TG | 3 | 0.47 ± 0.15 | 3 | 0.48 ± 0.17 | 0.93 |
| HDL-C | 3 | 0.79 ± 0.19 | 3 | 1.02 ± 0.29 | 0.24 |
| LDL-C | 3 | 0.78 ± 0.05 | 3 | 1.11 ± 0.36 | 0.13 |
| GLU | 3 | 4.45 ± 0.63 | 3 | 6.35 ± 1.97 | 0.13 |
| Ca | 3 | 2.49 ± 0.07 | 3 | 2.45 ± 0.80 | 0.59 |
| P | 3 | 2.02 ± 0.16 | 3 | 2.16 ± 0.65 | 0.39 |
| K | 3 | 5.68 ± 0.62 | 3 | 5.77 ± 1.79 | 0.90 |
| Na | 3 | 142.00 ± 0.82 | 3 | 142.00 ± 46.70 | 1.00 |
| CL | 3 | 100.80 ± 2.62 | 3 | 102.23 ± 32.73 | 0.54 |

**ALT**, alanine aminotransferase; **AST**, aspartate aminotransferase; **GGT**, gamma-glutamyl transpeptidase; **ALP,** alkaline phosphomonoesterase; **AMY**, amylase; **TP**, total proteins; **ALB**, albumin; **Mg**, magnesium; **TBIL**, total bilirubin; **TBA**, total bile acid; **$TCO_2$**, total $CO_2$; **UREE**, urea; **TC**, total cholesterol; **TG**, triglyeride; **HDL-C**, high density lipoprotein cholesterol; **LDL-C** low density lipoprotein cholesterol; **GLU**, glucose; **Ca**, calcium; **P**, phosphorus; **K**, potassium; **Na**, natrium; **Cl**, chlorine. Significance was established using Student's t-test. Differences were considered significant at *P<0.05.
